# AP-4 mediated ATG9A sorting underlies axonal and autophagosome biogenesis defects in a mouse model of AP-4 deficiency syndrome

**DOI:** 10.1101/235101

**Authors:** Davor Ivankovic, Guillermo López-Doménech, James Drew, Sharon A. Tooze, Josef T. Kittler

## Abstract

Adaptor protein (AP) complexes have critical roles in transmembrane protein sorting. AP-4 remains poorly understood in the brain despite its loss of function leading to a hereditary spastic paraplegia termed AP-4 deficiency syndrome. Here we demonstrate that knockout (KO) of AP-4 in a mouse model leads to thinning of the corpus callosum and ventricular enlargement, anatomical defects previously described in patients. At the cellular level, we find that AP-4 KO leads to defects in axonal extension and branching, in addition to aberrant distal swellings. Interestingly, we show that ATG9A, a key protein in autophagosome maturation, is critically dependent on AP-4 for its sorting from the *trans*-golgi network. Failure of AP-4 mediated ATG9A sorting results in its dramatic retention in the *trans*-golgi network *in vitro* and *in vivo* leading to a specific reduction of the axonal pool of ATG9A. As a result, autophagosome biogenesis is aberrant in the axon of AP-4 deficient neurons. The specific alteration to axonal integrity and axonal autophagosome maturation in AP-4 knockout neurons may underpin the pathology of AP-4 deficiency.

## Introduction

Adaptor protein (AP) complexes have roles in the selection of transmembrane proteins (cargo) for inclusion into vesicles. AP complexes interact with sorting motifs within the cytoplasmic facing tails of cargoes, leading to their specific enrichment at sites on donor membranes. Upon motif recognition and binding to cargoes, AP complexes recruit coat proteins which assemble to generate free vesicles (Bonifacino, 2014). Of the five members of the AP complex family, AP-1 and AP-2 are the best understood thusfar, functioning in clathrin-dependent sorting from the *trans*-golgi network (TGN) and endocyctosis from the plasma membrane respectively. Assembling as hetero-tetramers, AP complexes require the presence of all subunits for their function (Dell’Angelica et al., 1998; Hardies et al., 2015; Mitsunari et al., 2005). Mutations in genes encoding all subunits of AP-4 (ε; *AP4E1*, β4; *AP4B1,* μ4; *AP4M1* and σ4; *AP4S1*) have been identified as leading to a complex hereditary spastic paraplegia (HSP) termed AP-4 deficiency syndrome (henceforth: AP-4 deficiency) (Abou Jamra et al., 2011; Tesson et al., 2015). AP-4 deficiency patients present with early-onset severe intellectual disability, absence of speech and progressive spasticity leading to para- or tetraplegia (Abdollahpour et al., 2015). Anatomically, characteristic thinning of the corpus callosum and ventriculomegaly is evident in patients with AP-4 deficiency (Abdollahpour et al., 2015; Moreno-De-Luca et al., 2011; Verkerk et al., 2009). Despite this severe pathology little is known of AP-4 other than its localisation to the TGN and its clathrin-independence (Dell’Angelica et al., 1999; Hirst et al., 1999). The cargoes sorted by AP-4 in neurons and the functional consequence of their altered handling and subsequent trafficking as a result of disruption of the AP-4 complex remain poorly understood. Given the known roles of AP complexes in transmembrane protein sorting, identifying neuronal AP-4 cargoes will lead to a better understanding of the mechanisms underlying the pathology in AP-4 deficiency.

Macro-autophagy (henceforth: autophagy), the process by which organelles and macromolecules are recycled for the maintenance of cellular homeostasis can be simplified into three fundamental steps; induction, autophagosome biogenesis and lysosomal degradation. Progression through the autophagy pathway is in part mediated by the concerted recruitment of autophagy related (Atg) proteins (Mizushima et al., 2011), the sequence and necessity of which are conserved in the neuron (Maday and Holzbaur, 2014).

After the induction of autophagy, membrane elongation of sites on the endoplasmic reticulum (ER) forms a phagophore which incorporates cytosolic components (Ktistakis and Tooze, 2016). Enclosure of the expanding edges of the phagophore produces a double-membraned autophagosome, which then may fuse with late endosomes and lysosomes to form degradative autolysosomes (Galluzzi et al., 2017). Intact and efficient autophagy is of critical importance to post-mitotic neurons which cannot overcome proteotoxic burden through cellular division (Vijayan and Verstreken, 2017). The axon in particular represents a unique logistical challenge for autophagy due to its extreme length and architecture (Ariosa and Klionsky, 2015). Indeed, neurons have compartmentalised specialisation of autophagy; axonally derived autophagosomes exhibiting distinct maturation states from those generated somatodendritically (Maday and Holzbaur, 2016). Autophagosomes are constitutively generated in the distal axon (Maday et al., 2012), and are subsequently retrogradely trafficked toward the soma for their clearance by resident lysosomes in order to prevent distal accumulation (Xie et al., 2015). Thus, machineries necessary for autophagosome generation must be delivered to the distal axon to maintain effective biogenesis within this compartment. Given this, ATG9A is of particular interest as the sole mammalian transmembrane Atg identified to date, since it relies upon vesicular sorting and trafficking mechanisms for its distribution. Given the roles of ATG9A in phagophore extension and autophagosome maturation (Webber and Tooze, 2010a; Karanasios et al., 2016), its efficient sorting and subsequent delivery to the axon may be of critical importance for the maintenance of constitutive generation of autophagosomes in the distal axon. Intriguingly, in AP-4β null mice missorted AMPA receptors accumulated in autophagosomes in axonal swellings positive for LC3 (Matsuda et al., 2008). Whether a failure in autophagy underlies the pathology in AP-4 deficiency remains to be ascertained.

Here we identify neuroanatomical defects in an AP-4ε knockout mouse model mirroring those of AP-4 deficiency patients. Hippocampal neurons cultured from AP-4ε KO animals exhibit defects in axonal extension and branching, along with sites of distal swelling. We also show that functional AP-4 is critical for TGN exit of ATG9A, loss of which results in ATG9A retention within the TGN and reduction in axonal ATG9A, leading to aberrant autophagosome maturation in the distal axon. The impairment of axonal autophagosome biogenesis may underpin the severe pathology evident in AP-4 deficiency patients.

## Results

### AP-4ε ^(-/-)^ mice recapitulate characteristic anatomical defects of AP-4 deficiency

Given the stark anatomical features of AP-4 deficiency, we sought to characterise an AP-4ε^(-/-)^ mouse model to elucidate the AP-4 dependent mechanisms underpinning the pathology in this condition. Heterozygous mice carrying one copy of the targeting cassette (Fig 1A) were crossed, giving litters with AP-4ε^(+/+)^, AP-4ε^(-/-)^, and AP-4ε^(-/+)^ (hereafter WT, KO and HET respectively). KO was confirmed by PCR (Fig 1B) and AP-4s protein shown to be absent in KO embryos at E16 (Fig 1B). Brain regions of adult mice were also investigated, and AP-4ε shown to be absent at the protein level (Fig 1C). To examine whether loss of AP-4ε alters brain anatomy, sections were prepared from WT and KO animals at P30 and stained with NeuN and GFAP revealing brain morphology (Fig 1D). KO brains exhibited striking enlargement of the lateral ventricles at this timepoint (relative area: WT 1 ± 0.14, KO 10.12 ± 2.5, p = 0.0064; t-test). Staining axonal neurofilament-200 (NF200) (Fig 1F) revealed thinning of both the corpus callosum and dorsal fornix, axonal tracts projecting from the cortex and hippocampus respectively (Fig 1G; thickness corpus callosum: WT 181.1 ± 5.7 μm, KO 123 ± 6.9 μm, p = 0.0002. Fig 1H; thickness dorsal fornix: WT 119.2 ± 6.1 μm, KO 88.8 ± 8.9 μm, p = 0.0223; t-test). The identification of enlargement of the lateral ventricles and concurrent thinning of the corpus callosum are highly reminiscent of the characteristic features of AP-4 deficiency patients (Abdollahpour et al., 2015), supporting AP-4ε^(-/-)^ mice as a model of AP-4 deficiency.

**Figure 1:**
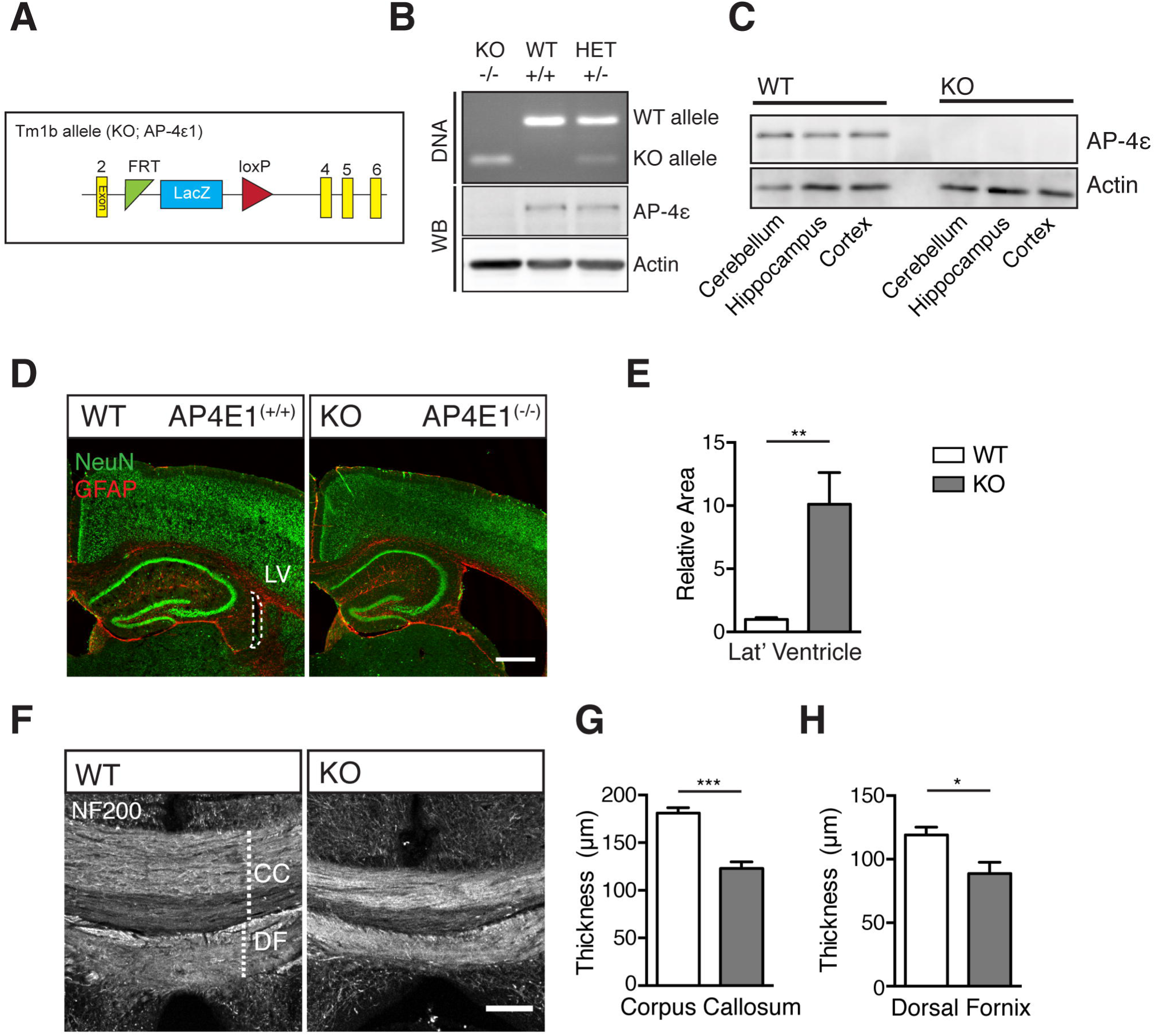
AP-4ε^(-/-)^ mice recapitulate characteristic anatomical defects of AP-4 deficiency. A. Schematic of the KO tm1b allele, showing removal of critical exon 3. B. Representative genotyping PCR of litter used for E16 hippocampal neuronal culture showing AP-4ε WT ^(+/+)^, KO ^(-/-)^ and Het ^(+/-)^ embryos. Bottom panel, western blot showing loss of ε protein in KO embryos. C. ε protein level in brain regions of adult mice (n = 3 repeats). D. E. Sections prepared from animals at P30 stained against NeuN and GFAP showing lateral ventricular enlargement in KO. Scale bar = 200 μm. (E) Quantification of relative area of lateral ventricle (n = 5 animals WT/KO). F - H. Commissural crossing axons, stained against Neurofilament-200 (NF200). Scale bar = 100 μm. (G) Quantification of thickness of CC and (H) DF (n = 5 animals WT/KO). Quantified data is expressed as mean ± SEM. (E) presented relative to control value (n = 5 animals per genotype). Statistical analysis: Two-tailed unpaired Student’s t-test, p<0.05, **P< 0.01 and ***P<0.001. CC - corpus callosum, DF - dorsal fornix.

### Axon specific defects in AP-4ε^(-/-)^ neurons

We next sought to establish whether defects at the cellular level were responsible for the thinning of the commissural axonal tracts in AP-4 KO animals. We examined GFP-transfected neurons at DIV-4 to evaluate the integrity of AP-4 KO axons (Fig 2A). KO axons exhibited reduced extension (Fig 2B; length: WT 963.5 ± 76.2 μm, KO 679.2 ± 51.9 μm, p = 0.0055; t-test) and branching (Fig 2C; number: WT 5.6 ± 3.2, KO 3.7 ± 1.9, p = 0.023; Mann-Whitney U test U test). Together these reductions likely underpin the thinning of axonal tracts in KO animals. In quantifying axonal length and branching parameters, we also noticed distal axonal swellings in KO neurons (Fig 2D-E; number: WT 1.5 ± 0.61, KO 7.2 ± 1, p < 0.0001; Mann-Whitney U test). Despite these alterations to the axon at this age in culture, nascent dendritic processes exhibited no alteration in total length (Fig 2F; length: WT 411.6 ± 37.6 μm, KO 384.5 ± 19.1 μm, p = 0.502; t-test), nor branching (Fig 2G; number: WT 10.41 ± 1.1, KO 10.04 ± 0.77, p = 0.772; t-test). Whether AP-4 KO had an impact upon developed dendrites was investigated by transfecting hippocampal neurons with GFP, revealing dendritic morphology at DIV-14 (Fig 2H). Investigation of the dendritic arbour using Sholl analysis did not reveal any alteration to the complexity of KO neurons (Fig 2I; NS between all concentric 10 μm regions; Two-way ANOVA), nor were any alterations in total dendritic length (Fig 2J; length: WT 2807 ± 326.1 μm, KO 2730 ± 324.2 μm, p = 0.87; t-test), nor branches per neuron (Fig 2K; number: WT 55.6 ± 4.9, KO 51.3 ± 5.1, p = 0.56; t-test) found. Notably we did not observe any dendritic swellings, despite axonal swellings still being evident at this timepoint in KO neurons (Fig S1; per 100 μm: WT 0.23 ± 0.04, KO 1.1 ± 0.2, p = 0.005; t-test). Together these parameters highlight the specific alteration in the integrity of axons in AP-4 KO neurons, whereas there was no alteration to dendritic complexity nor integrity.

**Figure 2:**
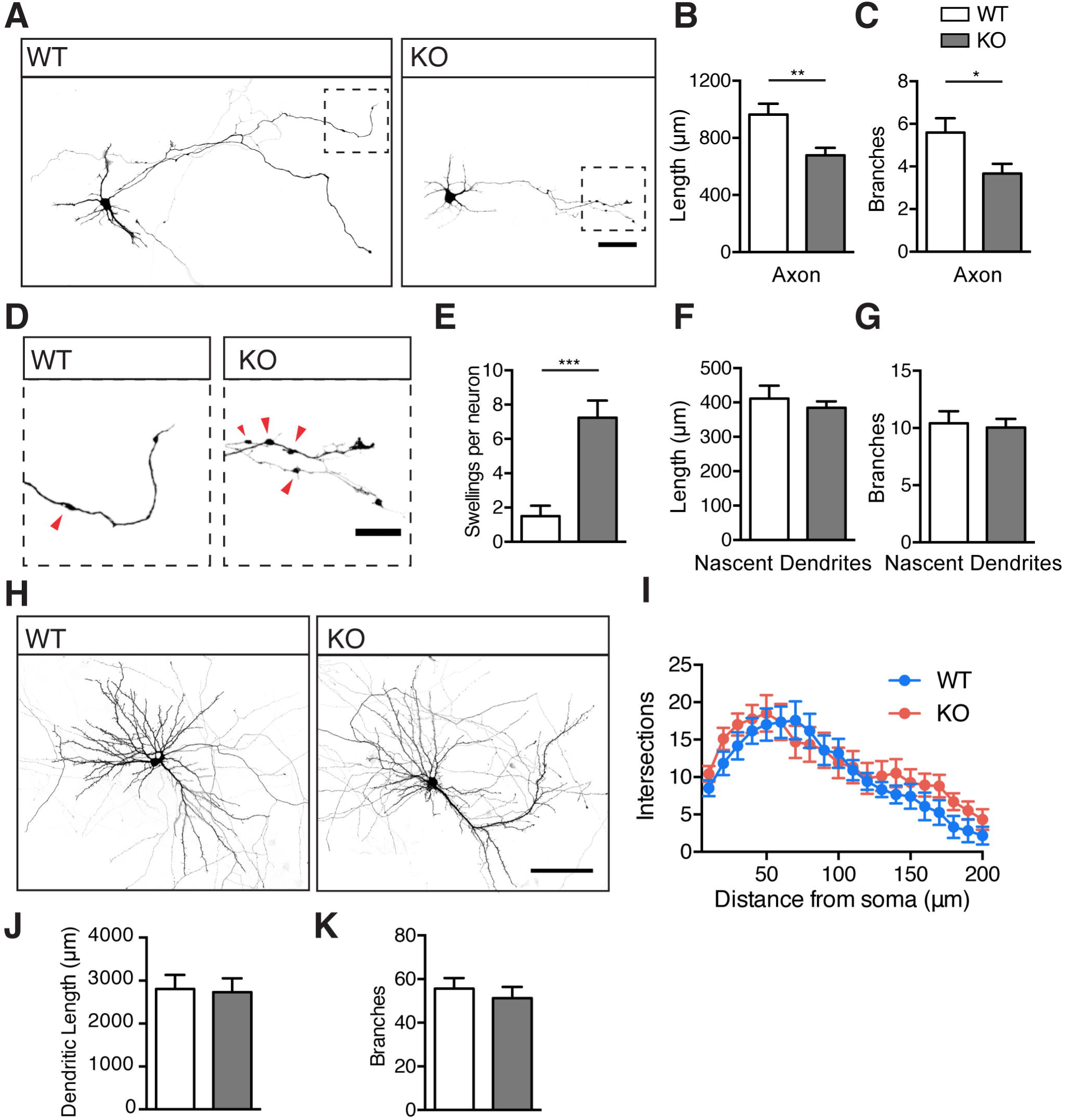
Axon specific defects in AP-4ε^(-/-)^neurons. A - C. Cultured GFP-filled DIV-4 hippocampal neurons stained against GFP showing neuronal morphology. Scale bar = 50 μm. (B) Quantification of total axonal length and (C) branches (n = 22/18 neurons WT/KO). D, E. Inset magnified panel from (A) of distal axonal regions, red arrows indicating axonal swelling. Scale bar = 20μm. (E) Quantification of number of swellings per neuron (n = 20/17 neurons WT/KO). F, G. Quantification of total nascent dendritic length (F) and branches per neuron (G) at DIV-4. (n = 22/27 neurons WT/KO). H - K. Cultured GFP-filled DIV-14 hippocampal neurons stained against GFP showing neuronal morphology. Scale bar = 100 μm. (I) Analysis of dendritic complexity using 10 μm concentric sholl intersections. Quantification of (J) total dendritic length and (K) total branches per neuron (n = 12/17 neurons WT/KO). Quantified data is expressed as mean ± SEM, from three independent experimental repeats. Statistical analysis: (I) Two-way ANOVA with Bonferroni post-hoc test. (B, F, G, J, K) Two-tailed unpaired Student’s t-test. (C, E) Two-tailed Mann-Whitney U test test, *p<0.05, **P< 0.01 and ***P<0.001.

### ATG9A accumulates in AP-4ε KO neurons

Interestingly, axonal swellings reminiscent of those we describe here in AP-4ε KO neurons have been identified in autophagy deficient models (Hara et al., 2006; Nishiyama et al., 2007). Moreover, the transmembrane protein ATG9A was identified as a putative AP-4 interactor by mass spectroscopy (Mattera et al., 2015), suggesting that ATG9A trafficking or sorting may be altered in AP-4ε KO neurons. We confirmed biochemically the interaction between ATG9A and the AP-4 complex by co-IP of ATG9A and AP-4 from adult mouse brain (Fig 3A), suggesting that ATG9A is indeed an AP-4 cargo in the brain. In further support of this we found that ATG9A was increased at protein level in KO hippocampus at P30 (Fig 3B-C; relative protein: WT 1 ± 0.23, KO 2.33 ± 0.05, p = 0.0046; t-test), suggesting that ATG9A levels are affected by the loss of AP-4 function. In accordance with this, in sections prepared at P30 we found that ATG9A accumulated in AP-4 KO mice within distinct structures in neuronal cell layers (Fig 3D, S2), highlighting alteration to ATG9A localisation *in vivo*. To better understand and confirm this accumulation within KO neurons, ATG9A levels and localisation were examined in cultured hippocampal neurons. We found near 3-fold accumulation of ATG9A in KO neurons, and stark retention within a reticular structure in the soma (Fig 3E-F; relative signal: WT 1 ± 0.06, KO 2.8 ± 0.2, p < 0.0001; t-test). These findings provide evidence towards a critical role for AP-4 in ATG9A handling in neurons both *in vivo* and in culture.

**Figure 3:**
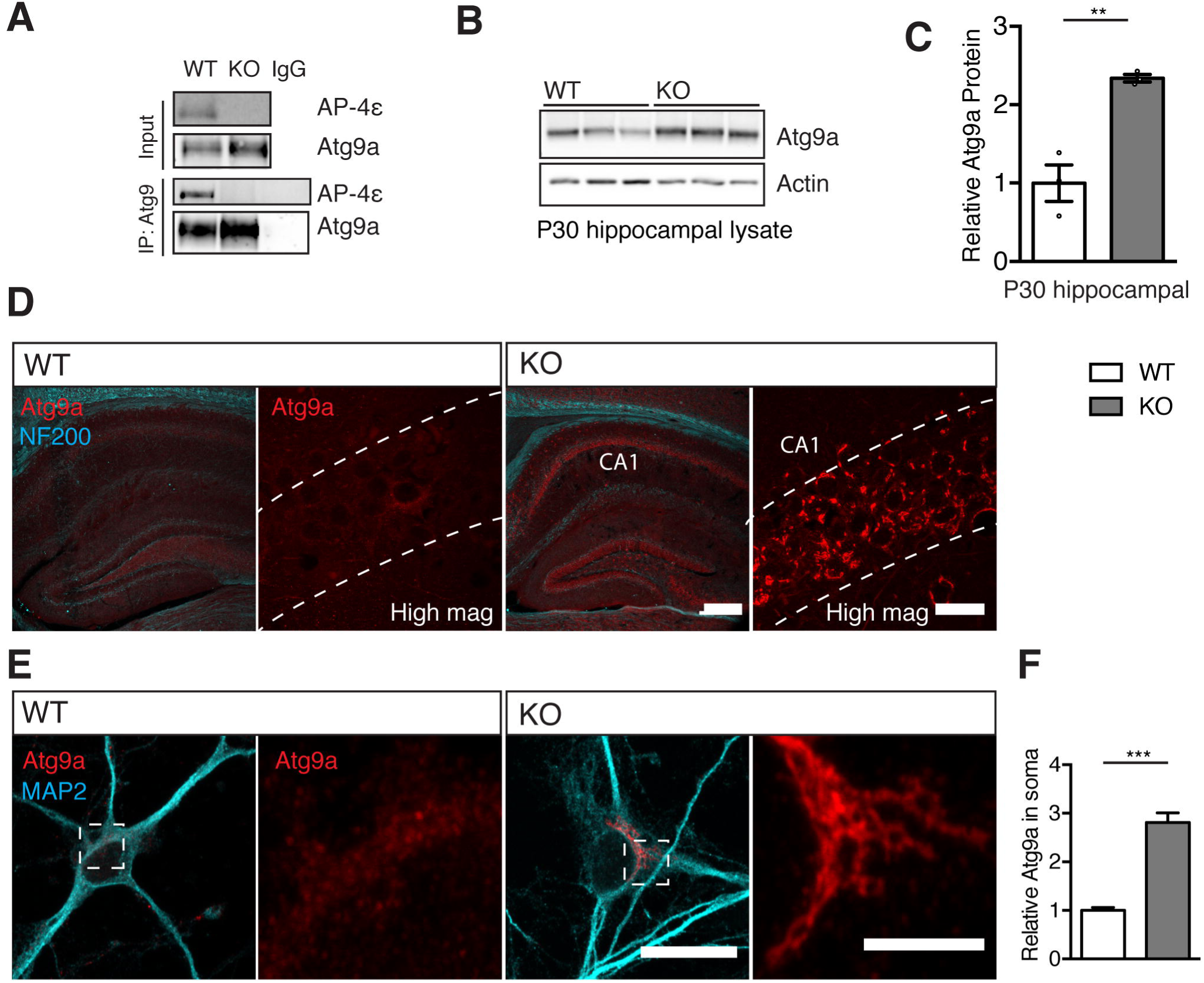
ATG9A accumulates in AP-4ε^(-/-)^ neurons. A, Endogenous co-immunoprecipitation of AP-4ε with ATG9A from mouse brain, showing interaction between AP-4 and ATG9A (n = 3 repeats). B, C. ATG9A protein accumulation in hippocampal lysates from KO animals and (C) densitometric quantification (n = 3 animals). D. Sections stained against ATG9A and NF200 revealing brain morphology and accumulation of ATG9A within cell layers of the hippocampus. High magnification panels show increased ATG9A immunoreactivity within the pyramidal CA1 cell layer. Scale bars = 200 μm, High mag 20 μm (n = 3 animals WT/KO). E, F. DIV-8 cultured hippocampal neurons stained against ATG9A and MAP2 revealing dendritic morphology. Inset panels show crops of cell body and accumulation of ATG9A. Scale bars = 20 μm, crop 5 μm (F) quantification of total ATG9A signal in neuronal soma (n = 40/20 neurons WT/KO). Quantified data is expressed as mean ± SEM, from three independent experimental repeats. Statistical analysis: Two-tailed unpaired Student’s t-test, **P< 0.01 and ***P<0.001.

### Functional AP-4 is critical for ATG9A exit from the TGN in neurons

The known functions of AP complexes in transmembrane protein sorting and AP-4 localisation to the TGN in cell lines lead us to investigate whether the somatic accumulation of ATG9A was due to its retention within the TGN as a result of the loss of function of AP-4. We firstly confirmed neuronal AP-4 localisation to the TGN and vesicles arising from it as evidenced by overlap of AP-4ε with the TGN marker Golgin-97 (Golg97) (Fig S3), supporting a role for AP-4 in transmembrane protein sorting from the TGN in neurons. We next sought to identify the compartment within which ATG9A was retained in culture and *in vivo* in AP-4s KO. Super-resolution structured-illumination imaging (SIM) of cultured neurons stained against *cis-golgi* (GM-130) and TGN (Golg97) markers revealed ATG9A to be highly associated with the TGN in KO neurons (Fig 4A-B), whereas in WT neurons ATG9A exhibited a vesicular localisation. The reticular ATG9A structures evident in KO neurons *in vivo* and in culture thus represent a pool of ATG9A that is retained within the TGN, indicating a critical role for AP-4 in ATG9A exit from the TGN.

**Figure 4:**
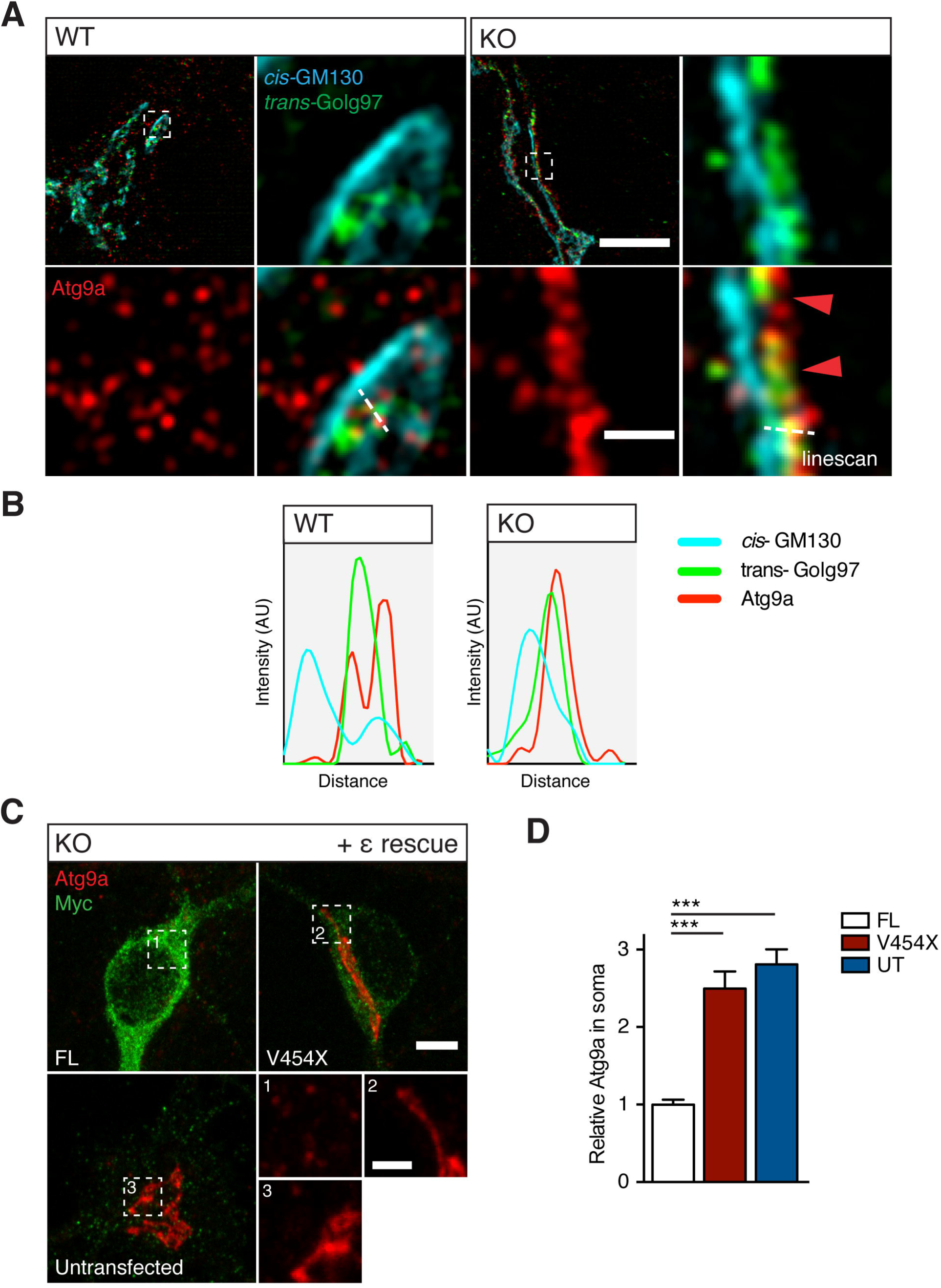
Functional AP-4 is critical for ATG9A exit from the TGN in neurons. A, B. SIM of DIV-8 cultured hippocampal neurons stained against ATG9A, *cis*-golgi marker GM130 and *trans*-golgi marker Golg97. Dashed boxes indicate region in magnified panels, showing vesicular ATG9A in WT neurons, and reticular ATG9A overlapping Golg97 in KO. (B) Intensity linescan demonstrates ATG9A retention within the TGN in KO neurons. Scale bars = 5 μm, 0.5 μm crop (n = 3 repeats). C, D. ATG9A constraint rescue by expression of myc-tagged FL and pathological V454x AP-4s constructs. Numbered dashed boxes indicate region in magnified panels. Scale bars = 5μm, crop 2 μm. (D) Quantification of relative ATG9A in soma of KO neurons in rescue conditions (n = 26/26/23 neurons FL/V454X/UT). (D) Quantified data is expressed as mean ± SEM, relative to FL rescue value from three independent experimental repeats. Statistical analysis: Kruskall-Wallis test, ***P<0.001.

We next sought to establish whether TGN ATG9A retention is a feature in AP-4 deficiency, through a rescue experiment using a pathological mutation identified in an AP-4 deficiency cohort (Fig S4) (Najmabadi et al., 2011). This homozygous 2-bp insertion resulting in frameshift and premature stop was identified in a consanguineous family with 3 of 4 children presenting with severe intellectual disability, microcephaly and spastic paraplegia. This mutation results in termination of AP-4ε within the trunk domain at V454 (Fig S4) which is likely necessary for correct AP complex assembly, stability and thus function (Peden et al., 2002). Given this, we hypothesised that the V454X-s pathological mutant would fail to rescue ATG9A localisation in AP-4 KO neurons. To this end, Myc-tagged full-length ε (FL-ε) and pathology associated V454X-ε were transfected in KO neurons, and ATG9A retention within the soma examined (Fig 4C, Fig S4). Restoration of ATG9A levels and vesicular localisation was evident with expression of FL-ε, whereas ATG9A remained TGN retained when V454X-ε was expressed, to a similar level as untransfected cells (Fig 4C-D; relative signal: FL 1 ± 0.06, V454X 2.48 ± 0.22, UT 2.81 ± 0.19, p<0.0001 V454X/UT to FL, NS between V454X and UT; Kruskall-Wallis). These findings highlight the necessity of functional AP-4 for ATG9A sorting, failure of which results in TGN ATG9A retention in AP-4 KO neurons and AP-4 deficiency.

### ATG9A TGN constraint results in defective axonal autophagosome maturation

Given the key roles of ATG9A in autophagosome biogenesis (Karanasios et al., 2016; Orsi et al., 2012; Webber and Tooze, 2010b) and overt axonal swellings reminiscent of autophagy deficient models in AP-4 KO axons, we hypothesised that autophagosome biogenesis is defective in KO neurons as a result of the impaired TGN exit of ATG9A. Indeed, we find that axonal delivery of ATG9A is reduced in KO neurons (Fig 5A, C; vesicles per 10 μm^2^: WT 5.56 ± 0.41, KO 3.49 ± 0.39, p = 0.0018; t-test), whereas dendritic ATG9A vesicle number is unaffected (Fig 5A, B; vesicles per 10 μm^2^: WT 4.47 ± 0.25, KO 4.18 ± 0.26, p = 0.43; t-test). The failure of AP-4 mediated ATG9A sorting from the TGN thus results in specific reduction of ATG9A trafficking to the axon, which may affect the capacity of axonal autophagosome biogenesis. To further investigate this we imaged the dynamics of LC3 (Fig 5D), which associates with autophagosomes from early through to late maturation states providing a robust marker for tracking autophagosomes throughout their lifespan (Maday and Holzbaur, 2014). Neurons were transfected with RFP-LC3 and movies captured within the distal most 100 μm of axon (Movies S1 and S2). Upon completion of biogenesis, autophagosomes generated in the distal axon must be trafficked retrogradely towards lysosomes resident in the soma for their degradation (Cheng et al., 2015). In KO axons, motile autophagosomes were found to exhibit reduced absolute retrograde displacement (Fig 5E; retrograde displacement: WT 10.43 ± 1.99 μm, KO 1.139 ± 1.42 μm, p < 0.0001; Mann-Whitney U test). Additionally, the mean retrograde run length per autophagosome was reduced, whereas anterograde run lengths were unaltered (Fig 5F; anterograde length: WT 15.89 ± 1.28 μm, KO 11.96 ± 1.16 μm, p = 0.686. retrograde length: WT 25.75 ± 1.59 μm, KO 12.58 ± 1.055 μm, p < 0.0001; Mann-Whitney U test). These analyses identified a specific reduction in the propensity of autophagosomes to move toward the soma in KO axons, suggesting altered maturation state. We also found that autophagosomes in KO axons were less motile, exhibiting reduction in total distance travelled (Fig 5G; length: WT 41.64 ± 2.11, KO 24.54 ± 1.72, p < 0.0001; Mann-Whitney U test). Indeed, KO autophagosomes spent more time stationary (Fig 5H % time: WT 58.75 ± 1.6 %, KO 65.82 ± 2.077 %, p = 0.0055; Mann-Whitney U test) and exclusively less time moving retrogradely (Fig 5H; % time retrograde: WT 23.45 ± 1.29 %, KO 15.52 ± 1.13 %, p = 0.0003. % time anterograde: WT 17.80 ± 1.17 %, KO 18.66 ± 1.55 %, p = 0.45; Mann-Whitney U test). Taken together, we identify autophagosomes to be less motile, and a specific reduction in the propensity to move retrogradely in distal KO axons. These findings suggest altered kinetics of autophagosomes during biogenesis in the distal axon, in accordance with the specific reduction in axonal ATG9A and accumulation in the soma.

**Figure 5:**
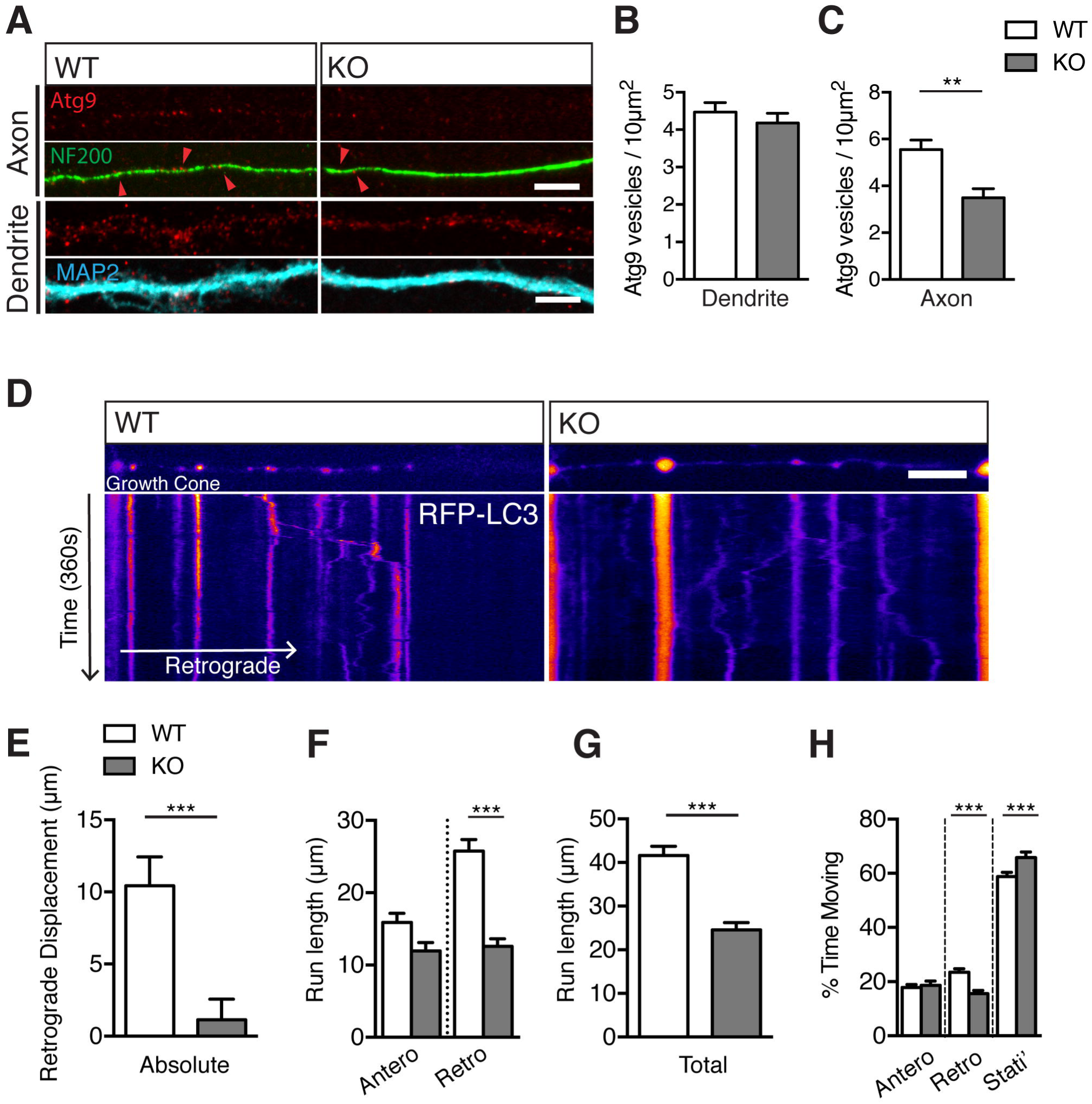
ATG9A constraint results in defective axonal autophagosome maturation. A - C. ATG9A vesicles in axons and dendrites. Quantification of vesicular density in (B) dendrites and (C) axons. Scale bar = 5 μm. (Dendrite; 28/24 WT/KO, Axon; n = 19/12 WT/KO). D - H. Live imaging of autophagosome motility at the growth cone and distal most 100 μm of axon. Movies generated over 6 minutes from cultured hippocampal neurons at DIV-6/7 transfected with RFP-LC3. First frames and resulting kymographs shown with pseudocolouring of RFP-LC3 signal. X-axis scale bar = 10 μm, Y-axis represents time (1px/1.5s). Quantification of; (E) absolute retrograde displacement (F) anterograde and retrograde run length per motile autophagosome, (G) total distance travelled per motile autophagosome, (H) proportion of time spent stationary, or moving anterogradely or retrogradely per motile autophagosome. (n = 227/117 motile autophagosomes from 46/36 neurons WT/KO). Quantified data is expressed as mean ± SEM, from three independent experimental repeats. Statistical analysis: (B, C) Two-tailed unpaired Student’s t-test. (E, F, G, H) Two-tailed Mann-Whitney U test, **P< 0.01 and ***P<0.001.

## Discussion

Despite the mounting evidence of impaired autophagy in neurodevelopmental and neurodegenerative disease (Alvarez-Erviti et al., 2010; Lee et al., 2011; Nixon et al., 2005; Shibata et al., 2006; Winslow et al., 2010; Xie et al., 2015), mechanistically autophagy remains poorly understood in the neuron. Neurons have a limited capacity to upregulate autophagy (Maday and Holzbaur, 2016), and as such may be particularly vulnerable to impaired autophagic flux. Given the spatial restriction of autophagosome biogenesis and the extreme length of the axon, it is critical that the delivery of newly synthesized cargoes from the soma is tightly balanced with efficient clearance to prevent accumulation at the distal axon. In the present study, we identify a critical role of AP-4 in axonal autophagosome biogenesis through the sorting of ATG9A from the TGN. Defective axonal delivery of ATG9A where AP-4 function is lost results in aberrant autophagosome maturation in the distal axon which may underpin the defects in axonal integrity in AP-4 deficiency. These findings strengthen the emerging links between HSP and autophagy (Chang et al., 2014; Khundadze et al., 2013; Oz-Levi et al., 2012; Vantaggiato et al., 2013; Varga et al., 2015), highlighting the importance of autophagy in the development and maintenance of axonal integrity.

Autophagosome generation is constitutive within the axon, predominantly occurring within the distal-most regions and at presynaptic sites (Maday and Holzbaur, 2014; Okerlund et al., 2016). Maturing autophagosomes initially exhibit bidirectional movement (Maday et al., 2012) prior to switching to robust dynein driven retrograde movement mediated by JIP1 (Fu et al., 2014). Interestingly, we found that a functional consequence of the reduction in the axonal delivery of ATG9A in AP-4ε KO neurons were alterations to autophagosome kinetics in the distal axon. Autophagosomes in KO axons were not only less motile, but also spent proportionally less time moving retrogradely and exhibited reduced net retrograde displacement. This specific reduction in the propensity of autophagosomes to move retrogradely in KO axons supports impaired autophagosome maturation in this compartment, in accordance with the specific reduction in the provision of ATG9A. Indeed, in *C.elegans* where AP-4 is not evolutionarily conserved (Boehm and Bonifacino, 2002), axonal delivery of ATG9A is also critical for axonal autophagosome biogenesis and axon outgrowth (Stavoe et al., 2016). It is also of note that in AP-4β deficient neurons AMPA receptors are missorted to axons, where they co-localise with LC3 positive autophagosomal accumulations through altered TARP (transmembrane AMPA receptor regulatory protein) dependent sorting (Matsuda et al., 2008). Given our identification of defective axonal autophagosome biogenesis in AP-4ε KO neurons, these likely accumulate as a result of the impaired autophagosome clearance due to ATG9A TGN retention. Whilst we cannot entirely rule out whether another unidentified AP-4 cargo contributes to alterations in autophagosome generation evident in the axon, the known roles of ATG9A make it a prime candidate in our AP-4 deficiency model. Indeed, in agreement with this ATG9A loss leads to axon swellings and thin corpus callosum in a CNS-specific knockout mouse model (Yamaguchi et al. 2017).

The distal axonal swelling evident in AP-4ε KO neurons have also been identified in studies where components critical for the early stages of autophagosome biogenesis are ablated, namely ATG5 and ATG7 (Nishiyama et al. 2007; Komatsu et al. 2006). Given that autophagosomes mature from ER sites (Ktistakis and Tooze, 2016; Maday and Holzbaur, 2014), it is intriguing that accumulation of expanded ER and ‘autophagosome-like’ structures are observed in the swellings of ATG5 null axons (Nishiyama et al., 2007) and that ER expansion is evident where ATG5 or Beclin-1 are silenced (Khaminets et al., 2015). Notably Atlastin-1, REEP1 and spastin, accounting for over 50% of HSP, all have roles in ER shaping and remodeling (Botzolakis et al., 2011; Montenegro et al., 2012; Park et al., 2010; Renvoisé and Blackstone, 2010), and axonal swellings have also been identified in several HSP models (Fassier et al., 2013; Tarrade et al., 2006; Watanabe et al., 2013). Whether axonal swellings comprise expanded ER as a result of impaired axonal autophagosome biogenesis remains to be elucidated. In the present study, our results are consistent with axonal swellings arising as a result of slowed or stalled autophagosome maturation in the axons of AP-4ε KO neurons.

70% of complex HSP patients presenting with progressive spasticity, intellectual impairment and thin corpus callosum are accounted for by mutations in SPG11 and SPG15, encoding spatacsin and spastizin respectively (Tesson et al., 2015). Notably, both spatacsin and spastizin have roles in autophagosome maturation and endolysosomal function (Vantaggiato et al., 2013). More recently, thin corpus callosum has been identified in autophagy-related CNS specific KO mouse models including ULK1/ULK2 dKO (Wang et al., 2017; Yamaguchi et al., 2017). Importantly, axonal extension is reduced in cultured primary neurons prepared from all of these lines (Khundadze et al., 2013; Pérez-Brangulí et al., 2014; Yamaguchi et al., 2017; Zhou et al., 2007). Given this similarity to AP-4ε KO mice in thin corpus callosum and concomitant reduction in axonal extension, we propose that thinning of the corpus callosum in AP-4 deficiency is due to the axonal extension defect elicited by failure of AP-4 mediated ATG9A sorting. Notably, constitutive knockout of both ATG9A and ULK1 leads to peri-natal lethality (Cheong et al., 2014; Kojima et al., 2015; Saitoh et al., 2009). Thus in AP-4 deficiency, the remaining pools of ATG9A that are not TGN retained likely account for the survival of AP-4 deficiency patients and AP-4ε KO mice into adulthood. We speculate that the dramatic increase in ATG9A within the TGN membrane leads to its stochastic incorporation into vesicles emanating from the TGN mediated by other carriers. As a result, sufficient vesicular ATG9A is delivered somatodendritically in KO neurons to maintain effective autophagy within this compartment. In accordance with this there is no defect in dendritic integrity in AP-4ε KO neurons despite the overt axonal defects evident. Alternatively, dendritic delivery of ATG9A may be mediated by another carrier, which would also account for the maintenance of the dendritic ATG9A pool. Specific axonal exclusion of somatodendritically destined ATG9A vesicles at the peri-axonal exclusion zone (PAEZ) (Farías et al., 2015) or the trafficking challenge posed by the axon may account for the reduced provision of ATG9A to the distal axon in this scenario.

During the preparation of this manuscript, we became aware of a study identifying AP-4 mediated TGN sorting of ATG9A in mouse fibroblast and HEK cell lines resulting in slowed autophagosome biogenesis (Mattera et al., 2017). The additional logistical challenge imposed by the great length of the axon would make autophagosome generation more critically dependent on AP-4 mediated TGN exit of ATG9A for neurons. This work further supports our findings *in vitro* and *in vivo*, lending weight to the implication of the failure of AP-4 mediated ATG9A sorting in our mouse model of AP-4 deficiency.

In summary, the present study reveals a critical role of AP-4 in sorting ATG9A from the TGN in neurons. Impairment of this function as evident in our AP-4 deficiency model leads to accumulation of ATG9A within the TGN *in vivo* and *in vitro,* leading to a specific reduction to the axonal delivery of ATG9A. Resulting defective axonal autophagosome biogenesis likely underlies the axonal defects evident in AP-4ε KO mice, providing evidence towards a mechanism of pathology in AP-4 deficiency.

## Materials and Methods

### Animals

AP-4ε knockout *(AP4E1^-/-^* C57BL/6J ^[Ap4e1tm1b(KOMP)Wtsi]^, KO) were generated using the knockout-first tm1a allele system (Skarnes et al., 2011) by the International Mouse Phenotyping Consortium (IMPC) MRC Harwell. Animals were maintained under controlled 12:12 hour light-dark cycles at a temperature of 20 ± 2°C with food and water ad libitum. Genotyping was carried out using the following primers; AP4E1-5arm-WTF: GCCTCTGTTTAGTTTGCGATG, AP4E1-Crit-WTR: CGTGCACAGACAGGTTTGAT and 5mut-R1: GAACTTCGGAATAGGAACTTCG. Littermate matched controls were used for primary neuronal cultures and immunocytochemistry experiments. All experimental procedures were in accordance with UCL institutional animal welfare guidelines, and under the UK Home Office licence in accordance with the Animals (Scientific Procedures) Act 1986.

### Antibodies and DNA Constructs

#### Antibodies

For immunocytochemistry (ICC), Immunohistochemistry (IHC) and western blotting (WB) antibodies were used with the following dilutions; Actin (Sigma A2066; WB: 1/1,000), AP-4ε (BD Biosciences 612018; WB: 1/300, ICC: 1/250), ATG9A (Rabbit STO-219 WB: 1/2000, IF: 1/2000 (Young et al., 2006)), ATG9A (Hamster 14F2 8B1; IF: 1/500 (Young et al., 2006)), GFP (Nescalai Tesque 04404-84; ICC: 1/1000), GFAP (Dako Z0334; ICC: 1/300), GM130 (BD Biosciences 610822; ICC: 1/1000), Golgin-97 (CST 13193; ICC: 1/250), MAP2 (Synaptic Systems 188-004, ICC: 1/500), NeuN (Chemicon MAB377; IHC: 1/300), NF200 (Abcam ab4680; IF: 1/500 IHC: 1/500) and MYC (NeuroMab 9E10; WB: 1/100, ICC: 1/100). HRP-Conjugated anti-mouse/rabbit antibodies were used for western blotting at 1/10,000 (Jackson Laboratories). Fluorescent Alexa Fluor conjugated secondary antibodies (Invitrogen and Abcam) for ICC, IHC and Super-resolution imaging were used as follows; anti-chicken 405 and 647, anti-Guinea Pig 405 and 647, anti-Mouse 488 and 647, anti-Rabbit 555 and 647. Anti-Armenian Hamster conjugated to Cy3 was used for super-resolution imaging (Jackson).

#### DNA Constructs

CAG-GFP (Addgene plasmid #16664), pmRFP-LC3 (Atkin et al., 2012)(Addgene plasmid #21075). Full-length N-terminally Myc tagged s was generated by cloning the coding sequence of AP4E1 (Cusabio; CSB-CL890772HU, cDNA clone MGC:163338) into pRK6-Myc. V454X-ε was then generated by reverse mutagenesis methods replicating the reported 2 nucleotide insertion leading to frameshift induced premature stop at V454 (Najmabadi et al. 2011).

### Brain lysate preparation for Co-Immunoprecipitation

#### Lysate Preparation

Brains to be used for co-immunoprecipitation were removed from animals and homogenized in ice cold HEPES buffer (50mM HEPES, 0.5% triton x-100, 150mM NaCl, 1mM EDTA, 1mM PMSF, 50μl Antipain/Pepstatin/Leupeptin in ddH_2_O). Lysate was solubilised by rotation for 2 hours at 4°C prior to ultracentrifugation at 38,000rpm for 40 minutes. Protein content determined using a BCA assay kit (Promega).

#### Co-Immunoprecipitation

5mg of brain lysate was incubated with 1μg of antibody in HEPES buffer for 12h at 4°C with rotation, and 1 μg IgG control (rabbit) was incubated with WT brain lysate in tandem. Input samples were incubated in the same manner at all steps as immunoprecipitation samples. Protein A agarose beads (Generon) were added for 4 hours to IP samples, beads washed in HEPES buffer and suspended in protein sample buffer (150mM tris pH 8, 6% SDS, 300mM DTT, 30% glycerol, 0.3% bromophenol blue) and heated to 95°C for 7 minutes prior to SDS-Page and western blotting.

### Brain Lysate preparation, SDS-Page and Western Blotting

#### Lysate Preparation

Brains to be used for western blotting were removed from animals and snap frozen at −80°C. For preparation of lysates, brains were defrosted, relevant regions dissected and kept on ice throughout. Tissue was homogenized by sonication in lysis buffer (50mM HEPES, 0.5% Triton-X100, 150mM NaCl, 1mM EDTA, 1mM PMSF and Antipain, pepstatin and leupeptin), and debris pelleted at 38,000g for 10 minutes at 4°C. Lysate protein content was determined using a commercial BCA assay kit (Promeaga) and samples denatured for 7 minutes at 95°C in protein sample buffer. Samples were stored at −20°C or - 80°C.

#### SDS-Page and Western Blotting

20-40μg of protein of protein lysate was separated by SDS-PAGE using XCell Minicell II systems (Novex) and transferred onto nitrocellulose (GE healthcare) or 0.45μm pore PVDF (for LC3, GE Healthcare). Membranes were blocked in milk (4% non-fat milk powder, 0.05% Tween-20 in PBS) for 1 hour and incubated with primary antibodies at empirically determined dilutions as above overnight with agitation at 4°C. Membranes were then washed, secondary HRP-conjugated antibodies applied in milk at 1/10,000 and after a final washing steps bands visualised by application of ECL substrate (Luminata Crescendo, Millipore) and imaging using a CCD based system (Quant LAS 4000, GE Healthcare). Densitometric analysis was performed using FIJI software (NIH).

### Hippocampal neuronal culture and transient transfection

#### Hippocampal neuronal cultures

Hippocampal cultures from crosses of heterozygous AP-4ε animals were prepared from embryos at E16 as described previously (Davenport et al., 2017; López-Doménech et al., 2016; Vaccaro et al., 2017). Briefly, hippocampi were dissected in ice-cold HBSS (Gibco) supplemented with 10mM HEPES and incubated in 0.25% trypsin for 15 minutes prior to trituration. Dissociated neurons were seeded onto Poly-L-Lysine (0.5mg/ml in 0.1M borate buffer, pH 8) coated coverslips at a density of 30-50,000/cm^2^ in attachment medium (10% horse serum, 10mM sodium pyruvate, 0.6% glucose in MEM (Gibco). Attachment media was replaced the next day with Maintenance medium (2% B27, 2mM glutamax, 100μg/ml Penicillin/Streptomycin in Neurobasal (Gibco). 50% of the maintenance medium replaced every 4 days after the first week in culture to maintain cell health.

#### Transient transfection

Neurons were transfected using lipofectamine 2000 (ThermoFisher) according to manufacturers protocols, at an empirically determined ratio of lipofectamine to DNA per construct used (GFP 0.25μg, RFP-LC3 0.25μg and ε constructs 1μg per 2 coverslips, 1μl lipofectamine per coverslip). Neurons were left to express constructs for 2-3 days prior to further experimentation.

### Immunocytochemistry and immunohistochemistry

#### Immunocytochemistry (ICC)

Hippocampal cultures on coverslips were fixed prior to staining with 4% PFA with 4% sucrose in PBS for 7 minutes at RT. Post-fixation coverslips were washed in PBS and permeabilised for 10 minutes in blocking solution (1% BSA, 10% horse serum, 0.1% Triton-X100 in PBS). Primary antibodies were diluted in blocking solution at empirically determined dilutions and applied for 1 hour at RT in a dark humidified chamber. Coverslips were washed in PBS and fluorescent-conjugated secondary antibodies as listed above were used at a concentration of 1/1000 and applied for 1 hour at RT in a humidified chamber. Coverslips were mounted in ProLong Gold mounting medium (Invitrogen, P36930) and allowed to dry overnight at RT prior to imaging.

#### Immunohistochemistry (IHC)

Brains were removed from animals and fixed by immersion in 4% PFA for 24h at 4°C, cryoprotected in 30% Sucrose-PBS for 24h and frozen and stored at-80°C. Frozen brains were embedded into OCT compound and serially cryosectioned into 30μm sections in a Bright OTF-AS Cryostat (Bright Instruments) and stored at-20°C prior to staining in cryoprotective solution (30% Glycerol, 30% PEG in PBS). IHC staining was performed with free-floating sections at RT with gentle agitation. Sections were washed and permeabilised in PBS-Tx (0.5% Triton-X100 in PBS) for 30 minutes prior to blocking in IHC blocking solution (3% BSA, 10% Fetal Bovine Serum, 0.2M Glycine in PBS-Tx) for 3 hours. A second block was applied for 3 hours as prior but with the addition of goat anti-mouse Fab-fragment (Jackson Immunoresearch) at 50μg/ml to reduced endogenous background when using antibodies raised in mouse. Sections were washed for 30 minutes and primary antibodies applied at concentrations as listed above in IHC blocking solution for 4 hours. Sections were washed for 30 minutes and fluorescent antibodies applied for 4 hours prior to mounting onto glass slides with Mowiol (Calbiochem) medium. Slides were allowed to dry at RT overnight prior to imaging.

### Image Analysis

All imaging and image analysis techniques were performed blinded. All WT and KO embryos generated per genotype were used, and cell numbers kept consistent between embryos rather than genotypes (as a result of blinding at acquisition stage). Between 3 and 6 images were taken per condition and samples sizes kept consistent across experimental techniques. All microscopic imaging unless stated otherwise was using an upright Zeiss LSM700 upright confocal microscope. Images were digitally captured using Zen 2010 Software (Zeiss), using oil immersion objectives: 63x; 1.4 NA, 40x 1.3 NA and air objectives; 10x 0.3 NA, 5x 0.16 NA.

#### Brain measurements

Quantification of the thickness of the corpus callosum and dorsal fornix was performed manually using Fiji. At least 2 brain sections per animal were analysed and the mean measurement used as the representative value.

#### Axonal Length, Branching and swellings

GFP-filled neurons at DIV-4 were fixed and imaged using a 40x objective, and images stitched using ‘MosaicJ’ or ‘Pairwise stitching’ (Preibisch et al., 2009) plugins in FIJI as required. Entire lengths of axons including all branches was measured manually using FIJI, and branches quantified excluding any process below 20 μm. Axonal swellings were defined as a compartment >2x the width of the axon shaft, and numbers counted manually. For DIV 14 swellings quantification, fields of view were captured, total axonal length and numbers of swellings present within the field captured were counted to determine swellings per 100 μm of axon.

#### Dendritic morphology and complexity

DIV 14 GFP-filled neurons were fixed and imaged, and images stitched as previously described where necessary. Dendritic morphology was reconstructed using Neuronstudio (CNIC) and inbuilt analysis tools used to ascertain total dendritic length and branches as described previously (Norkett et al., 2016; Pathania et al., 2014). Sholl analysis of intersections was performed using the ‘Simple Neurite Tracer’ plugin in FIJI, with a sholl radius of 10 μm. Images were stitched where necessary.

#### Nascent dendritic length and branching

DIV-4 GFP filled neurons were reconstructed using Neuronstudio and total length and branches of nascent dendritic processes per neuron quantified using inbuilt tools.

#### ICC quantification of total fluorescent signal

For quantification of dendritic and axonal vesicle numbers, regions positive for compartment markers (MAP2 and NF200 respectively) were outlined manually per image, and values normalised to area. Total fluorescence (ATG9A in soma) was quantified by outlining the cell soma and measuring total fluorescence using inbuilt FIJI tools.

### Live Imaging and autophagosome motility analysis

#### Live Imaging of autophagosome maturation

For autophagosome maturation and motility experiments, cultured hippocampal neurons were transfected at DIV-4 with RFP-LC3 as described, to be imaged at DIV 6-7. Imaging was carried out under perfusion with ACSF (124mM NaCl_2_, 2.5mM CaCl_2_, 2.5mM KCl, 1mM MgCl_2_, 10mM D-Glucose, 25mM NaHCO_3_, 1mM NaHPO_4_) at 37°C with a flow-rate of 1-2ml/min and aerated (5% CO_2_, 95% O_2_) throughout. Growth-cones were identified and the RFP-LC3 signal in at least the distal most 100 μm of axon captured using a EM-CCD camera system (iXon, Andor technology) mounted to an Olympus microscope (BX60M) with a 60x objective, as described previously (Norkett et al., 2016). A mercury arc lamp with filtering provided excitation of the RFP fluorophore (Cairn Research). Images were acquired using MicroManager (Opensource, Micro-manager.org)(Edelstein et al., 2014) for 6 minutes at 1 frame every 1.5s.

#### Autophagosome motility analysis

Movies generated from distal axons used to generate kymographs using the ‘Multiple Kymograph’ plugin. Resulting kymographs represent autophagosome motion as time on the y axis (1.5s/px) and distance on the x(0.1333μm/px). Trajectories were manually tracked and analysed using an in-house MATLAB script. Briefly, the motion of an autophagosome is possible to define precisely by the positional change from co-ordinates x^1^/y^1^ to x^2^/y^2^. Calculating all of the individual trajectory changes for an individual autophagosome’s track we were able to ascertain; velocity, proportion of time spent moving, directionality etc. Per track, portions of time spent moving at less than 0.05μm/s were classed as stationary.

### Structured Illumination Imaging (SIM)

SIM was performed on a commercially developed Zeiss Elyra PS.1 inverted microscope using a Zeiss 63x oil objective lens (NA: 1.4) and pco.edge CMOS camera and ZEN Black software (Zeiss) as described previously (Davenport et al., 2017; Norkett et al., 2016). Images were captured using SIM paradigms (34-μm grating, three rotations and five lateral shifts) and processed using the SIM reconstruction module within ZEN Black with default theoretical PSF and other settings. Shifts between acquired channels were corrected for using 100nm Tetraspec fluorescent microspheres (Molecular Probes).

### Statistical Analysis

Results were analysed using Graphpad Prism 6 (Graphpad Software Inc). Data is presented as mean ± SEM. Where normalized, values are presented relative to the average of control values unless stated otherwise. Data was tested for normality prior to statistical testing, and appropriate statistical tests used. For differences between two groups statistical significance was determined using unpaired two-tailed Student’s t-tests when parametric. Two groups were tested using two-tailed Mann-Whitney U tests where at least one group was non-parametric. For three or more groups, statistical significance was determined by two-way ANOVAs with Bonferroni post-hoc testing where data was parametric. Kruskall-Wallis H tests were used for comparison of three or more groups where atleast one group was non-parametric. Significance is represented as; p* < 0.05, p** < 0.01 and p*** < 0.001.

## Acknowledgements

The authors would like to thank all members of the Kittler Lab for invaluable discussions and suggestions. We extend thanks to Lorena Arancibia-Carcamo for support with script design and analysis methodologies. This work was supported by grants from the Medical Research Council (MR/N025644/1) and ERC (Fuelling Synapses) to J.T.K.. D.I. and J.D. were on the UCL Clinical Neuroscience Program funded by a Brain Research Trust PhD Scholarship and MRC PhD studentship, respectively. We thank the UCL Super-resolution Facility (funded by the MRC Next Generation Optical Microscopy Initiative) and the MRC LMCB Light Microscopy staff for their contributions.

## Author Contributions

This study was conceived by D.I. and J.T.K. Experiments were designed D.I., G. L.D. and J.T.K. and performed and analysed by D.I. and G.L.D. Analysis scripts were developed by J.D. and D.I.. S.T. provided essential advice, tools and reagents. D.I. and J.T.K. wrote the paper.

## Supplementary Figure Legends

**Figure S1.**
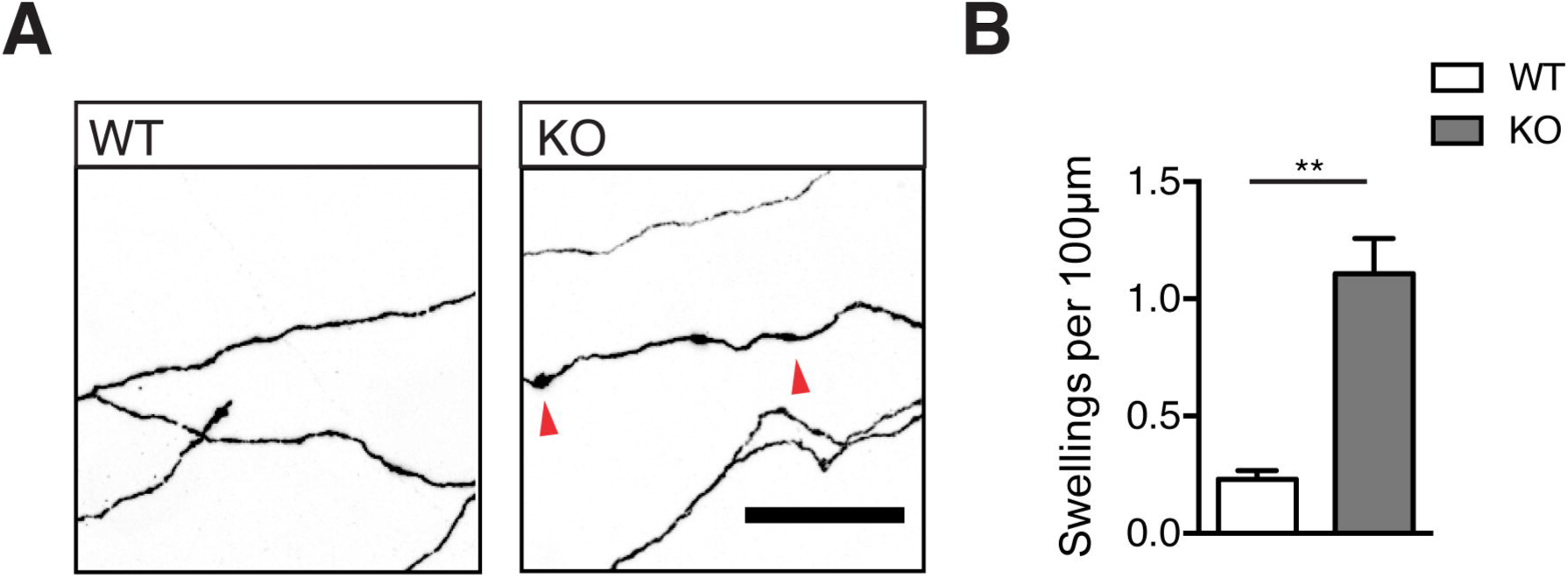
Axonal swellings are evident in DIV-14 hippocampal cultures. A, B. Axonal field of cultured GFP-filled DIV-14 hippocampal neurons stained against GFP showing axonal morphology. Scale bar = 20 μm (B) Quantification of axonal swellings per 100 μm. (n = 3/5 embryos WT/KO). Quantified data expressed as mean ± SEM, from three independent experimental repeats. Statistical analysis: Two-tailed unpaired Student’s t-test, **P< 0.01.

**Figure S2.**
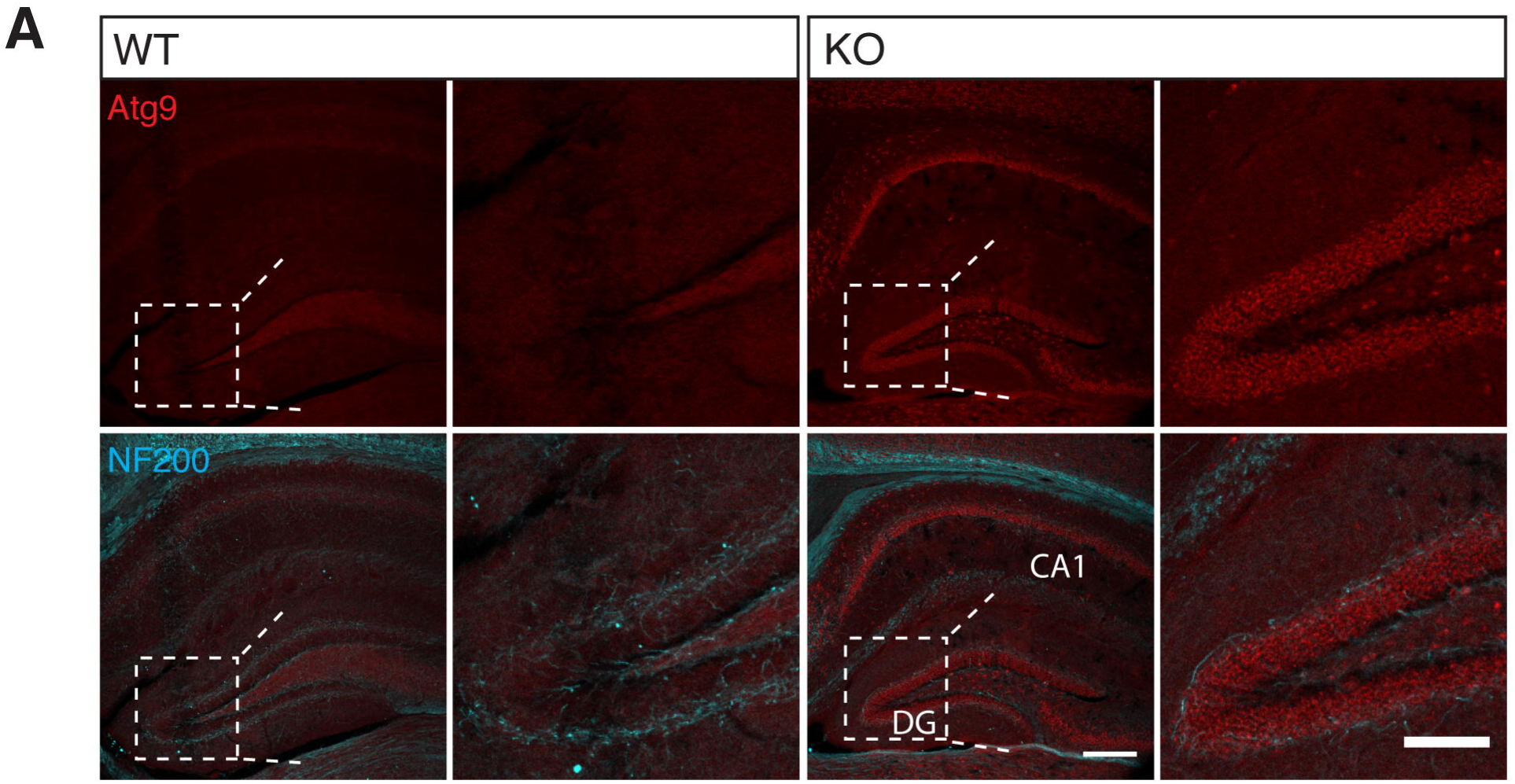
ATG9A accumulates in pyramidal cell layers of the hippocampus. A. Sections stained against ATG9A and NF200 revealing brain morphology and accumulation of ATG9A within cell layers of the hippocampus at P30 in AP-4 KO. High magnification panels show increased ATG9A immunoreactivity within the pyramidal dentate gyrus cell layer (n = 3 animals WT/KO).

**Figure S3.**
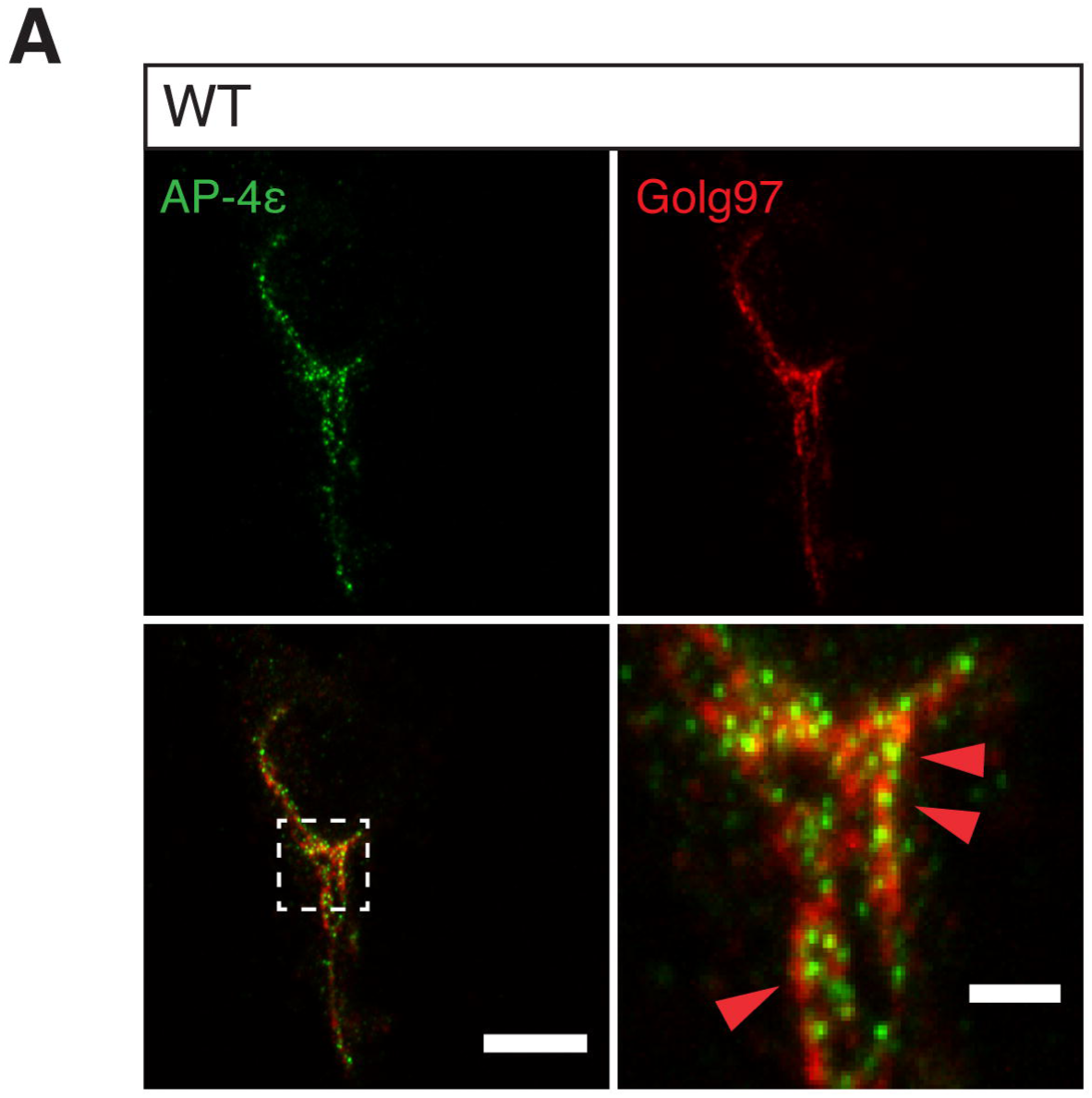
AP-4 is localised to the TGN in neurons. A. WT DIV-8 cultured hippocampal neuron stained against ε and Golgin-97 (Golg97). Dashed box shows magnified region and localisation of ε at the TGN. Scale bars = 10μm, 2μm crop (n = 3 repeats).

**Figure S4.**
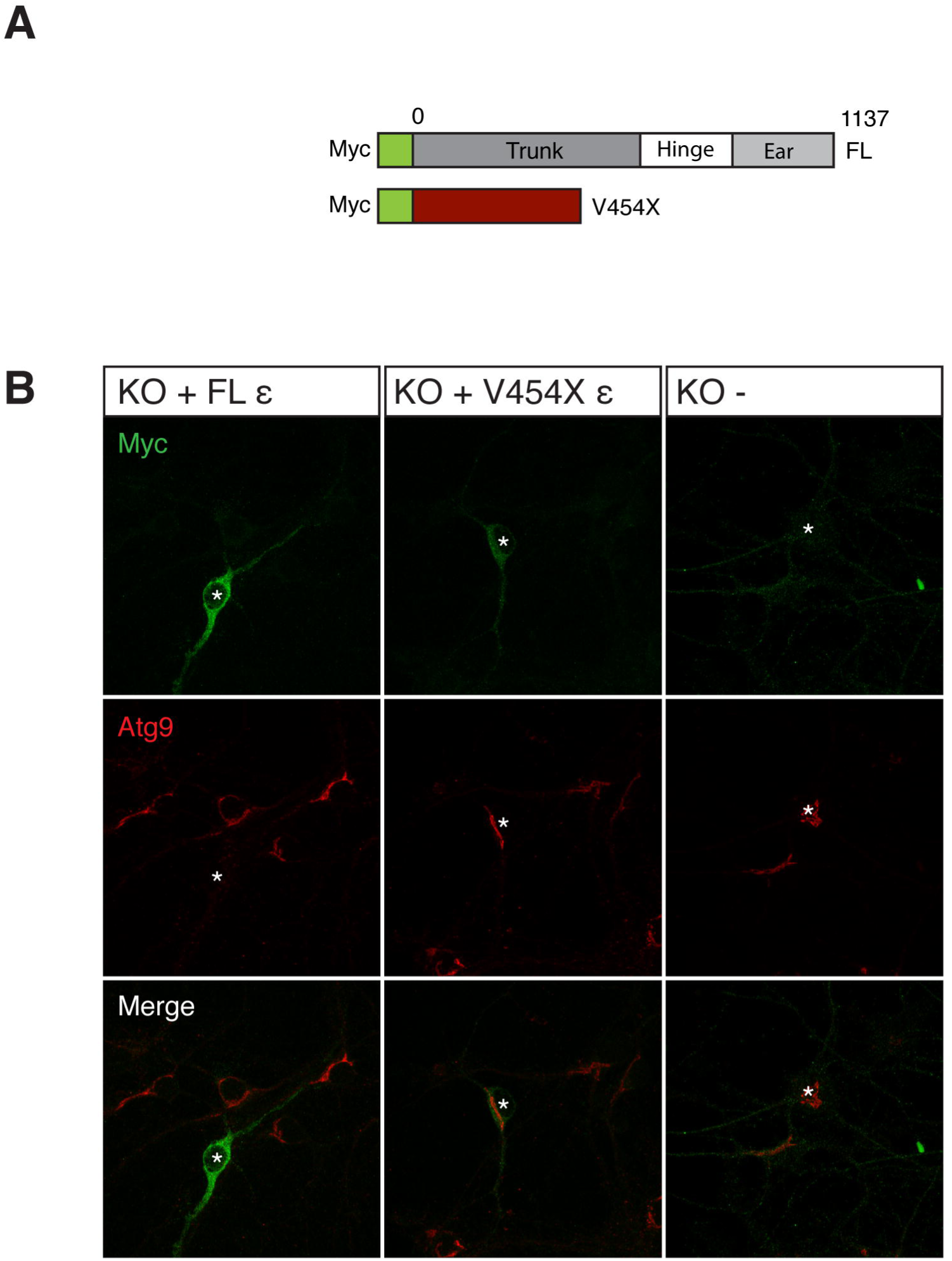
Reconstitution of AP-4 complex rescues ATG9A TGN constraint. A. Schematic of n-terminally myc tagged constructs generated for this study, showing structural position of pathological premature stop mutation. B. Related to figure 4C. Wide-field panels of exogenously expressed Myc constructs and endogenous ATG9A, showing rescue of TGN constraint only in transfected cells.

**Movies S1 and S2 – Defective distal axonal autophagosome maturation in AP-4 KO neurons**

Movies showing DIV 6-7 cultured hippocampal neurons transfected with RFP-LC3, pseudocoloured for clarity. Rightward motion is retrograde towards the soma, leftward is anterograde towards the growth cone. 1 frame/ 1.5s for 240 frames, playback at 20 frames/ second. S1 is representative movie from WT and S2 is representative movie from KO.

